# Genome-wide analysis of GATA factors in moso bamboo (*Phyllostachys edulis*) unveils that PeGATAs regulate shoot rapid-growth and rhizome development

**DOI:** 10.1101/744003

**Authors:** Taotao Wang, Yong Yang, Shuaitong Lou, Wei Wei, Zhixin Zhao, Chentao Lin, Liuyin Ma

## Abstract

**Background:** Moso bamboo is well-known for its rapid-growth shoots and widespread rhizomes. However, the regulatory genes of these two processes are largely unexplored. GATA factors regulate many developmental processes, but its role in plant height control and rhizome development remains unclear.

**Results:** Here, we found that bamboo GATA factors (PeGATAs) are involved in the growth regulation of bamboo shoots and rhizomes. Bioinformatics and evolutionary analysis showed that there are 31 PeGATA factors in bamboo, which can be divided into three subfamilies. Light, hormone, and stress-related *cis*-elements were found in the promoter region of the *PeGATA* genes. Gene expression of 12 *PeGATA* genes was regulated by phytohormone-GA but there was no correlation between auxin and *PeGATA* gene expression. More than 27 *PeGATA* genes were differentially expressed in different tissues of rhizomes, and almost all *PeGATAs* have dynamic gene expression level during the rapid-growth of bamboo shoots. These results indicate that *PeGATAs* regulate rhizome development and bamboo shoot growth partially via GA signaling pathway. In addition, *PeGATA26,* a rapid-growth negative regulatory candidate gene modulated by GA treatment, was overexpressed in Arabidopsis, and over-expression of *PeGATA26* significantly repressed Arabidopsis primary root length and plant height. The PeGATA26 overexpressing lines were also resistant to exogenous GA treatment, further emphasizing that PeGATA26 inhibits plant height from Arabidopsis to moso bamboo via GA signaling pathway.

**Conclusions:** Our results provide an insight into the function of GATA transcription factors in regulating shoot rapid-growth and rhizome development, and provide genetic resources for engineering plant height.

## Background

Moso bamboo is one of most-abundant non-timber forestry species and provides important resources for food, architecture, papermaking and fiber [1]. More importantly, moso bamboo is known for its explosive shoot growth rate, with a peak growth rate of 1 meter per day [1]. The rapid-growth shoot is largely dependent on the widespread rhizome system, which provide energy resources by absorbing from soil and more importantly, transporting from other rhizome-connected adult bamboos [2]. Therefore, studying the development of shoots and rhizomes will help us understand the rapid-growth regulation of bamboo and provide effective candidate genes for genetic manipulation of crop and forestry species.

The GATA factors play important roles in many developmental processes by binding to the consensus DNA sequence (A/T)GATA(A/G) to regulate gene expression at the transcriptional level [3, 4]. The GATA factors have a highly conserved type IV zinc finger DNA binding domain (CX_2_CX_17-20_CX_s_C) and followed by a basic region [5–7]. In animals, GATA factors typically contain two zinc finger domains (CX_2_CX_17-20_CX_2_C), while only the C-terminal domain has DNA binding function [5]. Animal GATAs are involved in development, differentiation, and control of cell proliferation [7]. However, the fungal GATA factors only contain a single zinc finger domain that is highly similar to the C-terminal zinc finger domain of the animal GATA factors [4, 8]. In plants, GATA factors contain CX_2_CX_18_CX_2_C or CX_2_CX_20_CX_2_C zinc finger domain [9, 10]. Interestingly, most of plant GATA factors have a single zinc finger domain, and very few of them also contain two zinc finger domains [9–11].

In animals, GATA factors involve in cell differential and organ development. Mutations in animal GATA factors cause severe developmental disorder diseases including anemia, deafness, renal and cardiac defects [12]. Fungal GATA factors play roles in nitrogen control, siderophore biosynthesis, light-regulated photomorphogenesis and circadian regulation [4].

Plant GATA factors originates from the identification of GATA motifs in regulatory regions of light and circadian clock responsive genes [13]. The first GATA factor identified in plant is NTL1 from *Nicotiana tabacum* [14]. GATA factors have been identified in many plant species, including Arabidopsis (29), rice (28), apple (35) and soybean (64) [9, 10, 15]. Plant GATA factors are involved in many developmental processes, including plant architecture [16], flowering development [17], hypocotyl elongation [18] and seed germination [19]. Plant GATA factors employ several underneath molecular mechanisms, such as modulate nitrogen metabolism [14, 20], act as transcriptional regulator by either integrity of light and phytohormone signal transduction [21, 22] or direct involvement in phytohormone signal transduction to regulate plant growth [23].

Plant GATA factors regulate light signal transduction by combining with GATA promoter of light related genes [24, 25]. *GATA2* (At2g45050) has also been identified as a key transcriptional regulator of the integration of light and brassinosteroid signaling pathways [22]. Recent evidences suggest that GATA factors are involved in the regulation of plant hormone signal transduction. Two orthologous GATA-type transcription factors- GNC and CGA1/GNL from Arabidopsis thaliana were identified as GA-regulated genes [21, 23]. Loss-of-function mutants and overexpression lines of *GNC* and *GNL* are functionally related to germination, greening, and flowering time [17]. Chromatin immunoprecipitation (CHIP) results show that these two genes are direct targets of PIF transcription factors, together with the fact that *gnc* and *gnl* loss-of-function mutations suppress *ga1* phenotypes, supporting that GNC and GNL are important repressors of GA signaling [21]. Another important phytohormone, auxin, is also regulated by GNC and GNL through functioning downstream of ARF2 [23]. In addition, the GATA factors are induced by cytokinin [26]. These results indicate that GATA factors play crucial roles in plant development and phytohormone-mediated growth. However, the role of GATA factors in rapid-growth and rhizome development remains elusive.

Recently, large-scale transcriptome analysis has shown that light and phytohormones may play important roles in the rapid-growth of bamboo [27–29]. In addition, a large number of transcription factor families are involved in the abiotic stress response and flower development have been studied in moso bamboo [30–32]. Although our group has functionally characterized rapid-growth associated key gene-*PeGSK1*, the rapid-growth associated transcription factor families are largely unexplored in moso bamboo.

In this study, we performed genome-wide survey of GATA factors in moso bamboo. A total of 31 GATA factors were identified in the moso bamboo genome. The phylogenetic relationship, gene structure and conserved domains of moso bamboo were systematically analyzed. The phytohormone-related *cis*-element and gene expression of *PeGATAs* under GA and auxin treatment were also characterized. More importantly, the gene expression of *PeGATAs* in different rhizome tissues and rapid-growth shoot were detailed analyzed. In addition, one of growth related PeGATA-PeGATA26 was overexpressed in Arabidopsis to functional validate its role in regulating plant height. Overall, our results provide information on the involvement of GATA factors in rhizome tissue development and rapid-growth shoot.

## Results

### Genome-wide characterization of GATA factors in moso bamboo

To identify the GATA factors in moso bamboo, the bamboo reference genome was used to scan the GATA factors using HMMER and blast tools (http://forestry.fafu.edu.cn/db/PhePacBio/download.php) [33]. A total of 31 potential GATA factors were identified in moso bamboo and named PeGATA1 to PeGATA31 based on the chromosomal location. The CDS and protein sequences of PeGATA genes were listed in Additional file 1 and 2: Table S1 and S2. The detailed information of these PeGATA factors including length of CDS, size of amino acid, molecular weight (MW) of protein, gene location on chromosome and isoelectric point (PI) were listed in Table 1.

**Table 1.** GATA factors in moso bamboo

| Name | Gene ID | Location | ORF length(bp) | Size (aa) | MW (kDa) | PI | Sub-family |
| --- | --- | --- | --- | --- | --- | --- | --- |
| PeGATA1 | PH01000001G0820 | 603239-605821(– stand) | 762 | 253 | 27.2 | 7.11 | □ |
| PeGATA2 | PH01000036G1110 | 651571-655167(+ stand) | 1212 | 403 | 42.4 | 5.2 | □ |
| PeGATA3 | PH01000040G1560 | 1013911-1015300(– stand) | 801 | 266 | 28.7 | 9.77 | • |
| PeGATA4 | PH01000114G0660 | 460195-464981(+ stand) | 912 | 303 | 32.2 | 5.96 | • |
| PeGATA5 | PH01000157G0800 | 521887-523791(– stand) | 654 | 217 | 23.2 | 6.15 | □ |
| PeGATA6 | PH01000162G1360 | 945504-954318(+ stand) | 1500 | 499 | 56.4 | 9.4 | □ |
| PeGATA7 | PH01000232G0180 | 85809-87165(– stand) | 420 | 139 | 15.6 | 9.23 | • |
| PeGATA8 | PH01000242G0460 | 296415-297844(– stand) | 648 | 215 | 22.6 | 8.2 | • |
| PeGATA9 | PH01000263G0760 | 473691-475362(– stand) | 1095 | 364 | 37.5 | 7.72 | □ |
| PeGATA10 | PH01000284G0590 | 365850-367843(– stand) | 1131 | 376 | 39.3 | 5.89 | □ |
| PeGATA11 | PH01000417G1130 | 669097-674119(– stand) | 948 | 315 | 35.8 | 8.87 | □ |
| PeGATA12 | PH01000468G1050 | 681872-683301(+ stand) | 663 | 220 | 22.7 | 5.78 | • |
| PeGATA13 | PH01000604G0620 | 351426-352613(+ stand) | 399 | 132 | 14.6 | 9.36 | • |
| PeGATA14 | PH01000750G0690 | 435897-442169(+ stand) | 777 | 258 | 28.1 | 8.18 | • |
| PeGATA15 | PH01000836G0660 | 444348-452795(– stand) | 1413 | 470 | 51.6 | 8.57 | • |
| PeGATA16 | PH01000985G0260 | 141878-143100(+ stand) | 402 | 133 | 14.8 | 9.87 | • |
| PeGATA17 | PH01001002G0190 | 175975-177694(+ stand) | 831 | 276 | 29.4 | 8.98 | • |
| PeGATA18 | PH01001129G0380 | 296297-297943(+ stand) | 699 | 232 | 26 | 9.16 | • |
| PeGATA19 | PH01001155G0480 | 343290-344746(– stand) | 741 | 246 | 25.3 | 9.66 | □ |
| PeGATA20 | PH01001253G0390 | 263024-267886(– stand) | 516 | 171 | 18.9 | 9.95 | • |
| PeGATA21 | PH01001451G0450 | 270274-272851(– stand) | 930 | 309 | 32.6 | 8.68 | □ |
| PeGATA22 | PH01001557G0370 | 244289-248001(+ stand) | 594 | 197 | 20.7 | 9.14 | • |
| PeGATA23 | PH01001584G0350 | 297120-303918(– stand) | 1035 | 344 | 37.8 | 4.75 | • |
| PeGATA24 | PH01001907G0160 | 109783-111580(+ stand) | 1017 | 338 | 36.2 | 9.26 | • |
| PeGATA25 | PH01002105G0190 | 151795-153666(+ stand) | 1248 | 415 | 44 | 8.6 | □ |
| PeGATA26 | PH01002473G0050 | 19960-21655(– stand) | 1011 | 336 | 36.1 | 9.64 | • |
| PeGATA27 | PH01002681G0110 | 59436-60698(– stand) | 690 | 229 | 24 | 8.49 | • |
| PeGATA28 | PH01002830G0260 | 174984-177126(– stand) | 381 | 126 | 13.8 | 9.69 | • |
| PeGATA29 | PH01003365G0100 | 53196-55342(+ stand) | 369 | 122 | 13.3 | 9.4 | • |
| PeGATA30 | PH01003433G0110 | 88789-92096(– stand) | 1167 | 388 | 39.8 | 9.33 | □ |
| PeGATA31 | PH01004789G0060 | 58264-61570(– stand) | 993 | 330 | 35 | 6.06 | □ |

bp: base pair, aa: amino acids, MW: molecular weight, PI: isoelectric point, kDa: kilodalton

The length of CDS ranges from 366 bp to 1,500 bp, and the length of proteins ranges from 122 aa to 499 aa (Table 1). PeGATA29 is the smallest GATA protein with 122 amino acids, and the largest protein is PeGATA6 with 499 amino acids (Table S1). The predicted molecular weight of 31 PeGATA proteins ranges from 13.3 kDa (PeGATA29) to 56.4 kDa (PeGATA6) with an average size of 29.86 kDa (Table 1). The predicted PI of 31 PeGATA factors are all below 10.0, and the minimal protein is PeGATA23 with only 4.75 (Table 1).

To further investigate and characterize sequence conservation in the GATA proteins, multiple sequence alignments were performed using the amino acid sequences of the conserved GATA motifs in 31 PeGATAs (Fig. 1). Most bamboo GATA factors contain a single zinc finger domain. However, unlike Arabidopsis, several bamboo GATA factors contain multiple zinc finger domains (Fig. 2). Most of bamboo GATA factors contain 18 residues in the zinc finger loop (CX_2_CX_18_CX_2_C), while five of them have 20 residues in the zinc finger loop (CX_2_CX_20_CX_2_C) (Fig. 1). Interestingly, the gene subfamily analysis revealed that all of these five PeGATA factors all belong to the Class C type of the PeGATA family (Fig. 3). Similar to Arabidopsis and rice, moso bamboo does not contain the animal- and fungal-type CX_2_CX_17_CX_2_C zinc finger domains (Fig. 1). Notably, five PeGATA genes factors have a defective GATA zinc finger domain (Fig. 1). PeGATA1 lacks the first Cys residue (-SHC) and PeGATA30 lacks the last Cys residue (CND-). Meanwhile, the GATA factors PeGATA14, PeGATA17 and PeGATA18 have only partial GATA motif (SRLTPAMRRGPTGPRSLCNAC for PeGATA14, CSDCNTTKTPLWRSGPCGPKAA for PeGATA17 and CSDCNTTKTPLWRSGP for PeGATA18) (Fig. 1). The observation is similar to the rice GATA factors as OsGATA24 also contains a partial GATA motif [9]. The results indicated that the bamboo GATA factors have a highly conserved GATA motif, especially compared to rice.

**Fig. 1.**
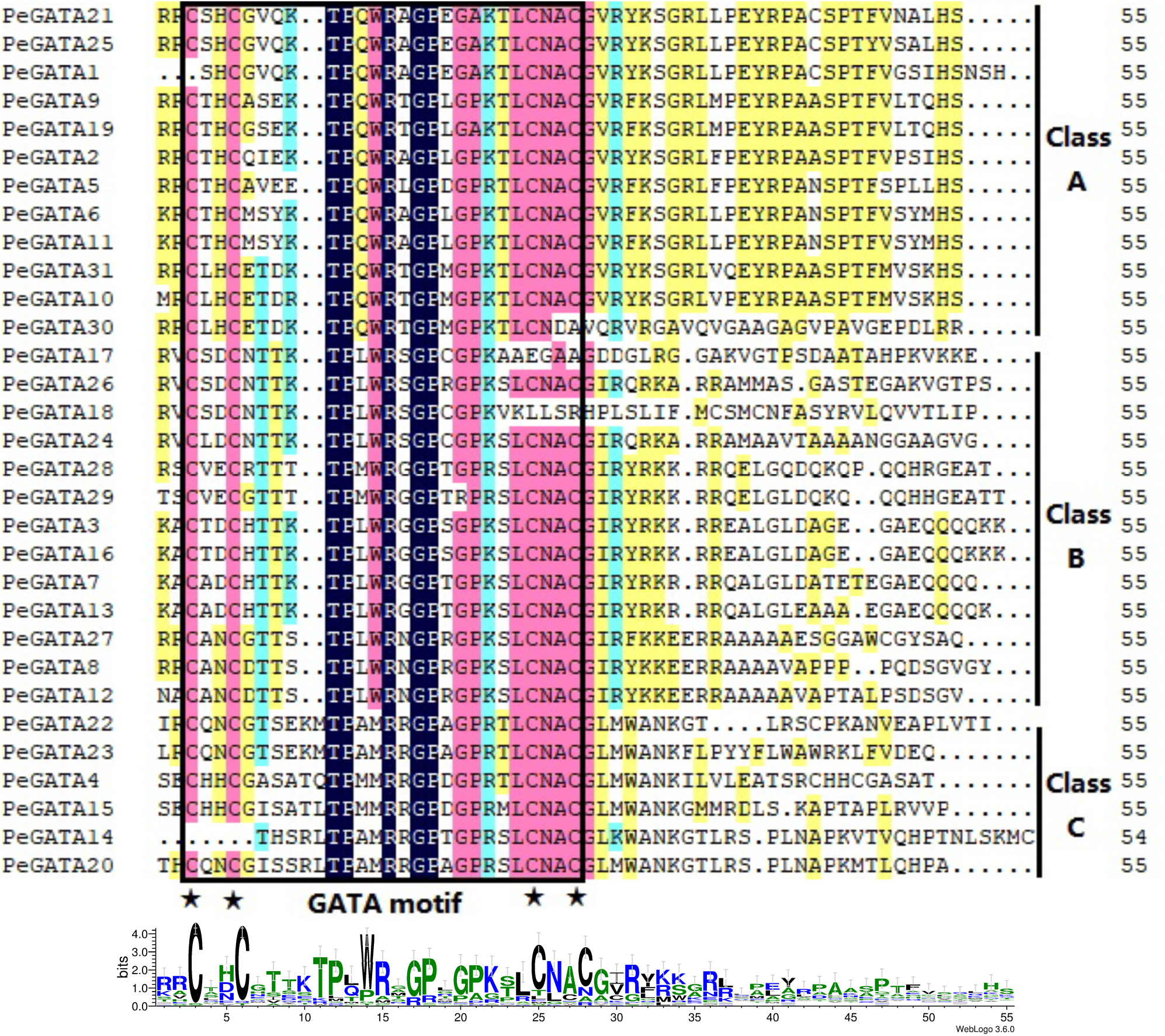
Alignment of the amino acid sequences of bamboo GATA factors. The GATA motifs and amino acid positions are marked with a box and an asterisk. The sequence identities of GATA motifs are shown at the bottom.

**Fig. 2.**
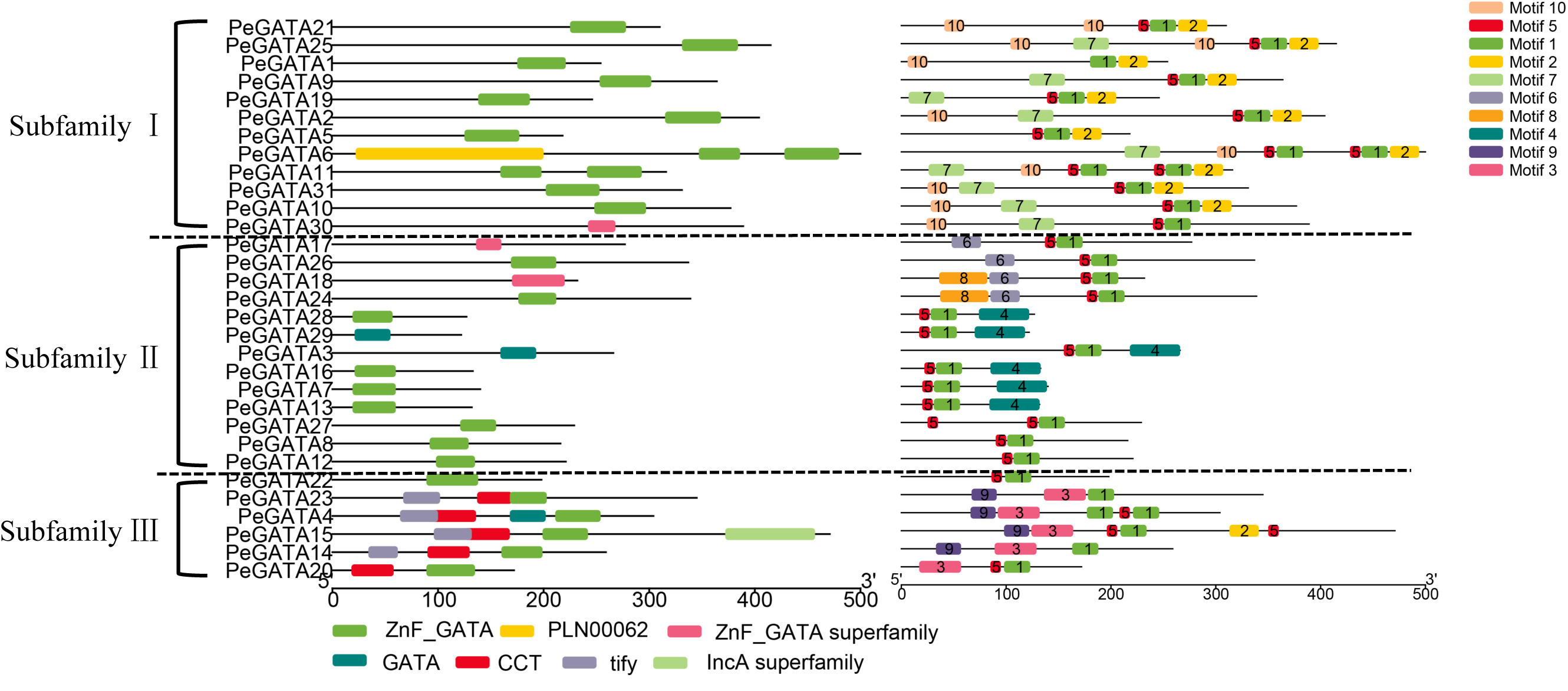
Schematic diagram of conserved domain analysis in bamboo GATA proteins. Each color represents a different motif.

**Fig. 3.**
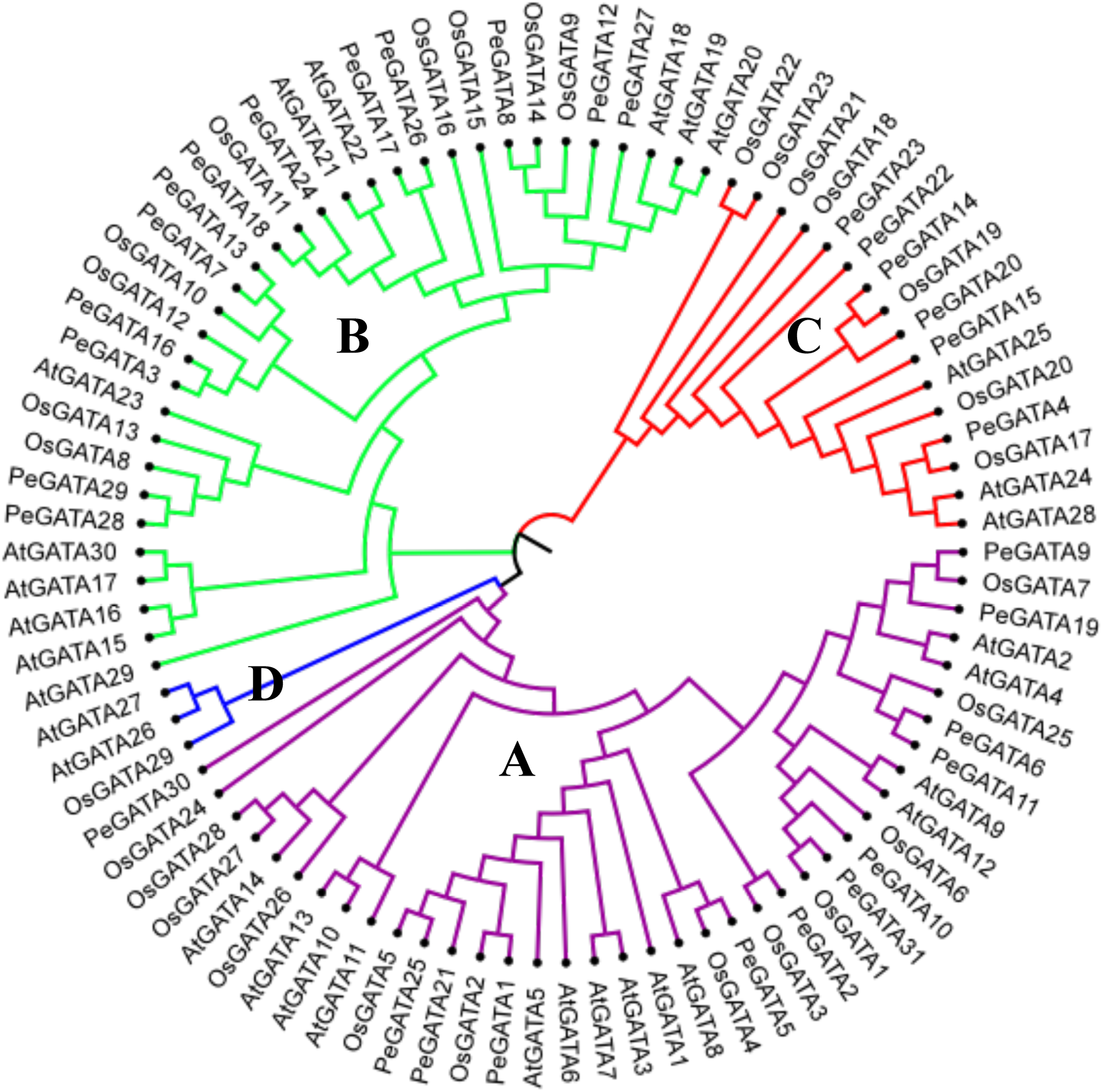
Phylogenetic analysis of GATA factors in bamboo, rice and Arabidopsis. The phylogenetic tree was made based on the amino acid sequences using MEGA7.0 by the neighbor-joining method with 1000 bootstrap replicates. The tree shows four major phylogenetic classes (Classes A to D) indicated by different colors.

To further reveal the diversification of GATA genes in moso bamboo, putative conserved functional domain and motifs were also predicted in the NCBI conserved domain database and program MEME. Through MEME analysis, 10 motifs among the different gene subfamilies is shown in Fig. 2 and the identified multilevel consensus sequence for the motifs is shown in Additional file 3: Table S3. Motif 1 and 5 presented in 29 PeGATA proteins and they were annotated as conserved GATA zinc finger domain CX_2_CX_18_CX_2_C and CX_2_CX_20_CX_2_C, respectively (Fig. 2). Motif 5 was not found in PeGATA1, PeGATA14 and PeGATA23 by MEME (Fig. 2), which may be attributed to the zinc finger GATA subfamily domain corresponding to the conserved domain. Motif 2, 7 and 10 appeared nearly all members in subfamily I, and motif 4 and motif 6 only appeared in subfamily II (Fig. 2). Motif 3 was identified as the CCT domain and motif 9 was identified as TIFY domain (Fig. 2). These two domains were specific to subfamily III that was consistent with the classification by conserved domain as shown in Fig. 2. The identification of subfamily-specific motifs from bamboo GATA factors suggests that these motifs may contribute to the functional differences among different subfamilies.

### Comparison analysis of the GATA subfamily among Arabidopsis, rice and moso bamboo

GATA factors in Arabidopsis and rice are classified in the clade A-D according to the residues of zinc fingers [9]. To determine the phylogenetic relationship among GATA genes in Arabidopsis, rice and moso bamboo, unrooted phylogenetic tree with 90 GATA factor sequences from all three species was constructed. The phylogenetic tree analysis shows that all GATA factors have three major clades (Classes A, B and C) (Fig. 3). Among them, Class A is the largest clade and contains 38 members. In this clade, twelve bamboo GATA factors (PeGATA1/2/5/6/9/10/11/19/21/25/30/31) clustered with the Arabidopsis GATA factors AtGATA1, AtGATA2, and AtGATA4, which have been reported to be involved in light regulation of gene expression and photomorphogenesis [22, 34]. Class B formed the second largest clade containing 33 members and 13 bamboo GATA factors (PeGATA3/7/8/12/13/16/17/18/24/26/27/28/29) clustered with the Arabidopsis GATA factors AtGATA21 (GNC) and AtGATA22. These two GATA factors regulates phytohormone response, chlorophyll biosynthesis, starch production, plant architecture, and nitrogen metabolism [17, 21, 23, 34, 35]. In Class C, six bamboo GATA factors (PeGATA4/14/15/20/22/23) clustered with the Arabidopsis GATA factor AtGATA25 (ZIM, Zinc-finger protein expressed in Inflorescence Meristem) and shows the potential roles of hypocotyl and petiole elongation [18]. It is worth noting that no bamboo GATA factor is found in Class D, which explains that bamboo GATA factors may have different functions compared to Arabidopsis and rice.

### Gene structure of bamboo *GATA* genes in moso bamboo

To determine the phylogenetic relationships among different members of the GATA factors in moso bamboo, a phylogenetic analysis based on alignments of the 31 full-length GATA protein sequences was performed. As shown in Fig. 1 and Fig. 3, the protein sequence alignment and neighbor-joining phylogenetic tree divides 31 PeGATAs into three clades according to the pattern of zinc finger domain or homologous domains to the Arabidopsis and rice GATA factor families. The gene structure of the *PeGATA* genes was shown in Fig. 4. The total exon numbers of *PeGATAs* from each subfamily were calculated. Subfamily I comprised 12 members with two or three exons except *PeGATA19* and *PeGATA6*. *PeGATA19* has only one exon and *PeGATA6* has more then three exons with long introns. Subfamily II consists of 13 members, and all of them contain two or three exons. Subfamily III was formed by included 6 members with five to twelve exons (Fig. 4). The gene structure of GATA factors is similar to that of rice [9]. Overall, the *PeGATA* genes contain exons ranging from one to twelve in its CDS, and the gene structure is obviously different from each other. The results indicated that the bamboo *GATA* genes have undergone significant changes during its long evolutionary history.

**Fig. 4.**
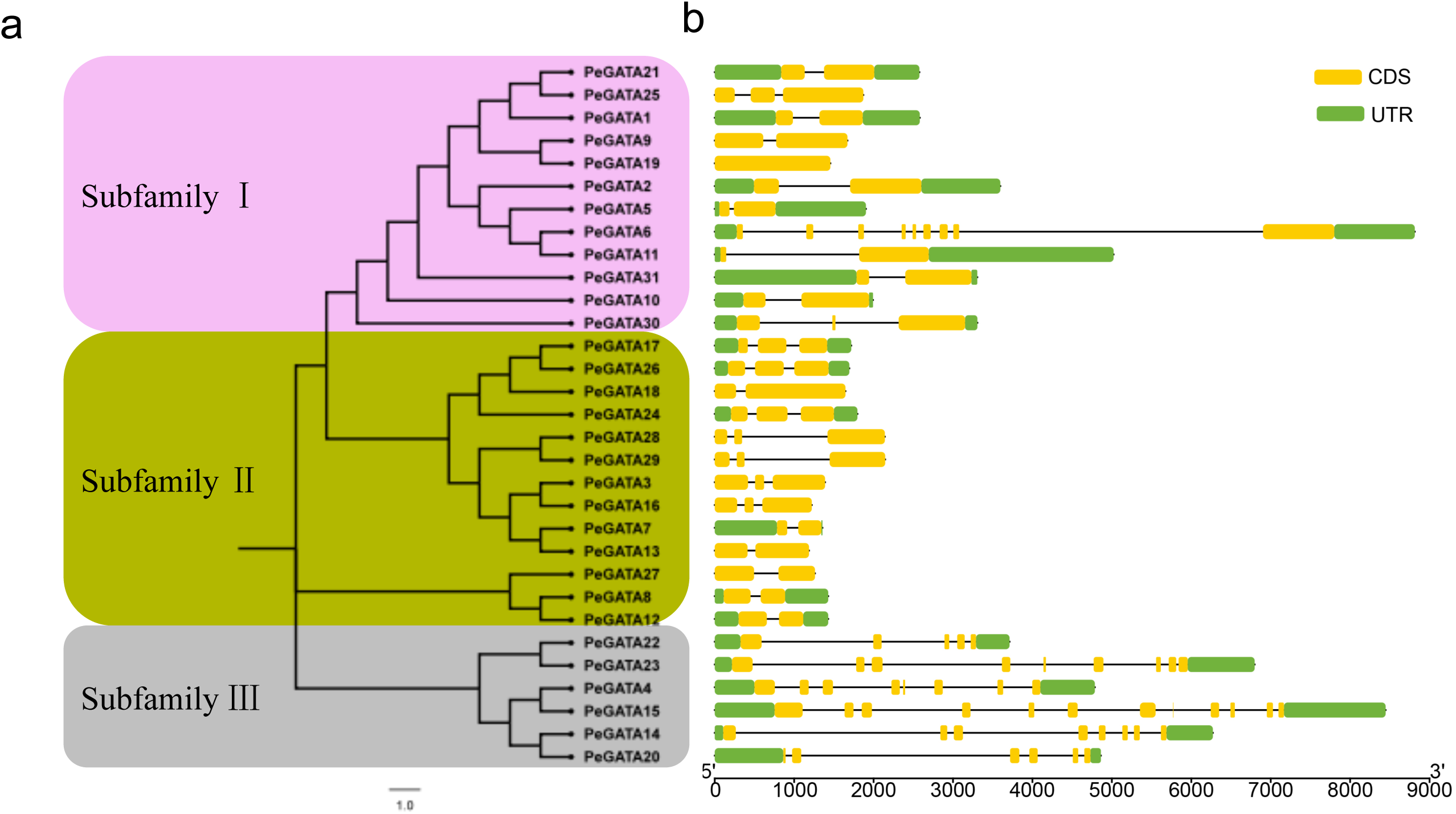
Phylogenetic analysis and gene structure of bamboo GATA factors. (a) Phylogenetic tree construction of the PeGATA factors based on the amino acid sequences using MEGA 7.0. The tree showed three major phylogenetic subfamilies (subfamilies I to ·), represented by different colored backgrounds. (b) CDS/UTR structure of the *PeGATA* genes. The yellow and green boxes indicate exons and UTRs, and the black lines represent introns. The size of exons and introns can be estimated using the scale at the bottom.

### Identification of hormone-related *cis*-elements in the promoter of the *PeGATA* genes

To further explore the function and regulatory pattern of the *PeGATA* gene, the PlantCARE database was used to scan the putative *cis*-elements inside the 1500 bp upstream of transcription start site. We categorized *cis*-elements into four categories based on their functions: light response elements, development, hormone and stress associated *cis*-elements (Fig. 5). The predicted *cis*-elements in *PeGATA* genes were closely related to the function of the GATA family in other plants [17, 19, 22, 23, 25]. Light responsive elements like G-box, GT1 and TCT were widely present in the promoter of *PeGATA* genes, and the G-box element has been reported to be involved in the regulation of chlorophyll II biosynthesis in Arabidopsis [36]. We also identified several hormone-responsive *cis*-elements such as ABRE [37], CGTCA-motif, TGACG, and TCA-elements (abscisic acid, MeJA and salicylic acid), and abiotic stress-responsive elements including ARE, GC-motif, LTR and MBS. In addition, tissue specific elements such as CAT-box, circadian responsive element and cell cycle regulation elements like MSA-like were also found in the promoter of the *PeGATA* genes, which may have function in the regulation of plant morphology, flowering and growth [38]. Overall, *cis*-elements analysis indicated that bamboo GATA factors might be involved in response to light and phytohormone to regulate growth. Transcription factors are typically located in the nucleus and regulate transcription of the target genes by binding to the *cis*-elements in their promoters. Consistent with our hypothesis, subcellular localization assays in tobacco showed that randomly selected bamboo *GATA* genes *PeGATA7, 20, 26* and *28* were clearly localized in the nucleus according to the GFP and DAPI stain signals (Fig. 6). Localization analysis revealed that bamboo GATA factors could also act as transcription factors to regulate gene expression.

**Fig. 5.**
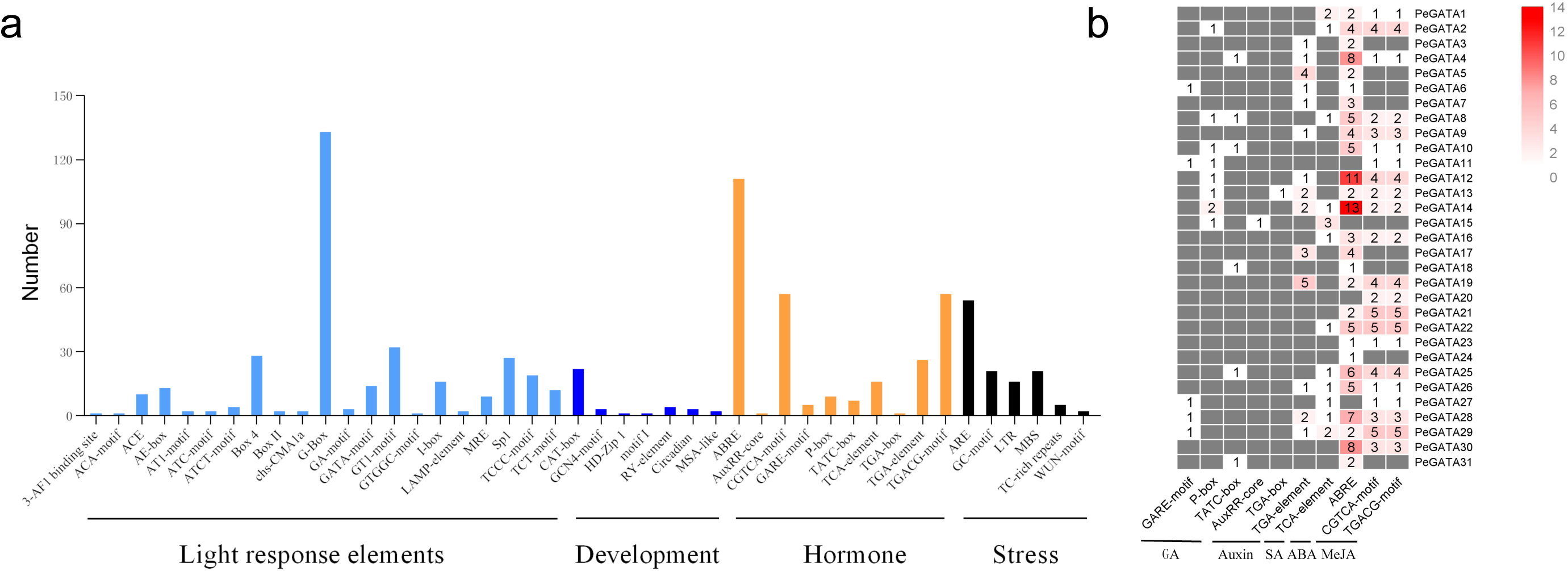
*Cis*-elements analysis in the promoter of bamboo *GATA* genes. (a): Overview of the main types of cis-elements identified from the 1.5-kb upstream sequence of the bamboo *GATA* genes by the PLANTCARE database. (b): Hormone related *c*A-elements were analyzed and each colored block with numbers represents the number of *cis*-elements in the bamboo GATA promoter.

**Fig. 6.**
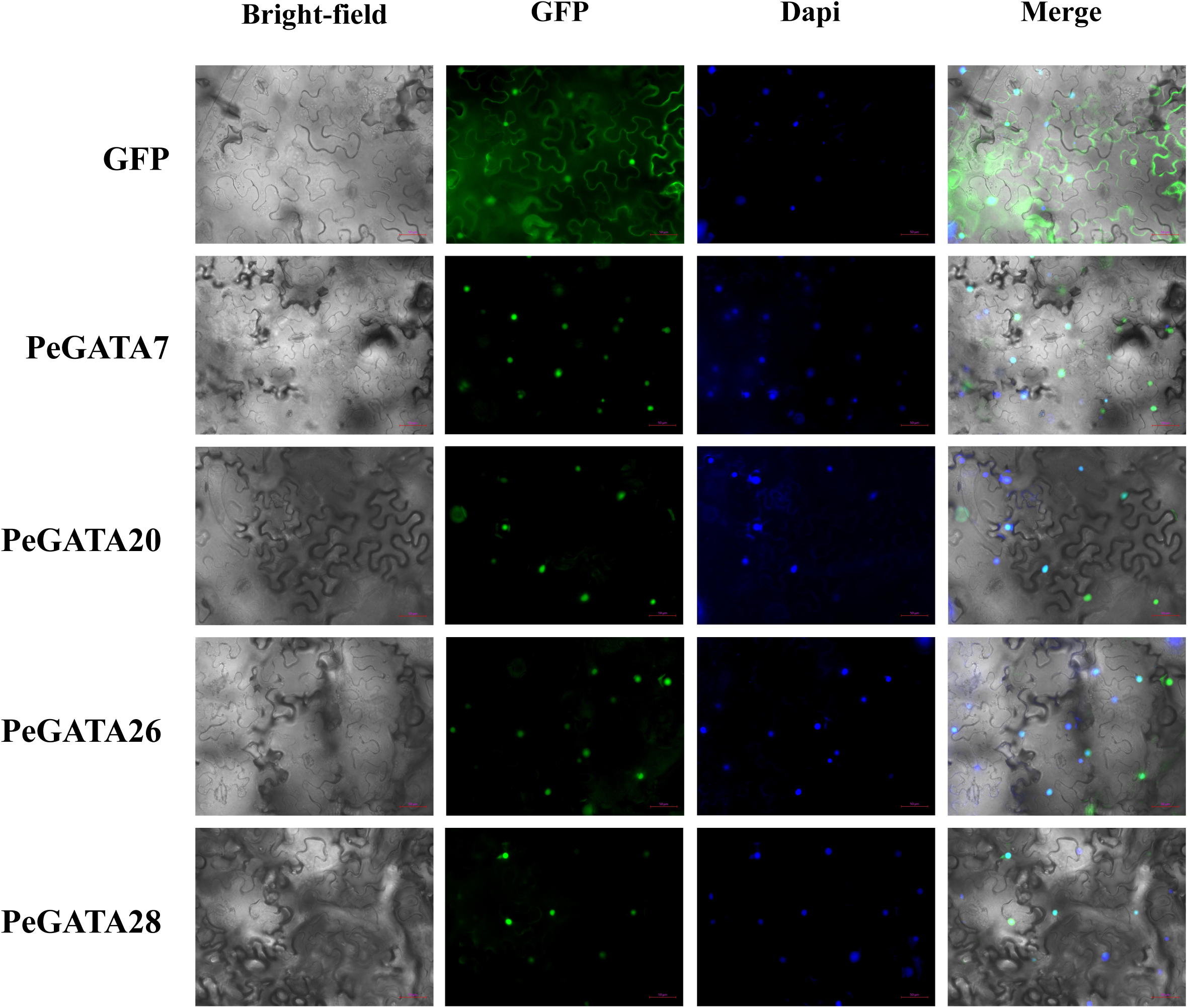
Subcellular localization analysis of bamboo GATA factors. The bamboo *GATA* genes were cloned and constructed in a modified pCambia3301 vector with a C-terminal GFP fusion. These vectors were transformed into tobacco, and the GFP and DAPI signals were captured from the identical areas by microscopy (20x).

### Dynamic gene expression pattern of *PeGATAs* in rhizome tissues

The bamboo rhizome system can be divided into three groups: lateral buds, rhizome tips, and new shoot tips [2]. The widespread rhizome system is essential for rapid-growth of bamboo shoot through adopting and utilizing nutrients including nitrate [39]. The GATA factors are also related to nitrogen metabolism in other species [20], so we firstly checked if GATA genes expressing differentially in lateral buds, rhizome tips, and new shoot tips. By analyze the RNA-seq data from our previous study [2], we showed significant differential expression pattern of *PeGATA* genes among different rhizome tissues (Fig. 7a, Additional file 4: Table S4). As shown in Fig. 7a, a total of 15 *PeGATA* genes (*PeGATA1, 5, 9, 10, 14, 18, 19, 20, 21, 23, 25, 26, 27, 28* and *29*) showed significantly higher expression in lateral buds than that from other two tissues. Five *PeGATA* genes (*PeGATA6, 7, 8, 11* and *22*) highly expressed in the new shoot tips, while reduced their expression in lateral buds. In the rhizome tips, seven *PeGATA* genes (*PeGATA2, 3, 4, 15, 16, 24* and *30*) have remarkable higher expression than other two tissues. In addition, *PeGATA12* and *31* expressed highly both in lateral buds and rhizome tips, while slightly expressed in new shoot tips. Overall, 29 of the 31 *PeGATA* genes showed differential expression in three bamboo rhizome tissues, suggesting that PeGATA factors may contribute to the growth regulation of rhizome.

**Fig. 7.**
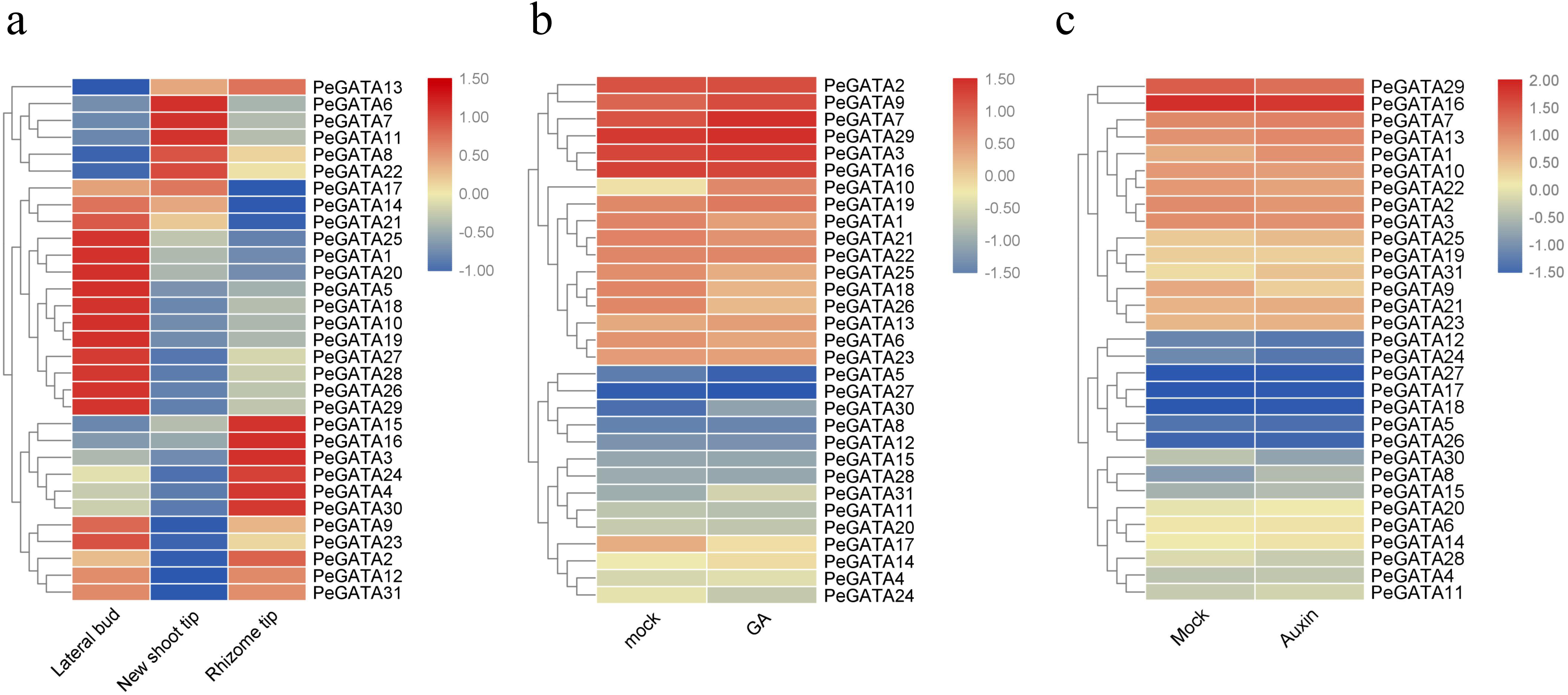
Expression profiles of bamboo *GATA* genes in different tissues and hormone treatment. (a): The gene expression of 31 bamboo *GATA* genes in different rhizome tissues was presented by heatmap. (b) and (c): The expression of bamboo GATA genes in the seedlings under GA and auxin treatment. Expression values were normalized and presented at the right side, and green represents lower expression level and red indicates a higher expression level.

### Expression profile of *PeGATAs* in bamboo under the treatment of exogenous phytohormone

GATA factors are closely related to phytohormones to regulate Arabidopsis growth and development [21, 23], take together with the identification of phytohormone related *cis*-elements in the promoter of the bamboo *GATA* genes (Fig. 5b), we rationally hypothesized that the *PeGATA* genes are also tightly regulated by phytohormones. To test our hypothesis, we performed gene expression analysis of the *PeGATA* genes under GA and auxin treatment based on the RNA-seq data published in the previous studies [28, 40]. A total of 12 *PeGATA* genes showed significant gene expression under GA treatment (Fig. 7b, Additional file 5: Table S5). Among them, the expression of *PeGATA7, 9, 10, 19, 30* and *31* was increased in GA_3_ (100 μΜ) treated seedlings compared to that from untreated control (Fig. 7b). The largest difference was observed in *PeGATA10* (increased by 3.22-fold after GA_3_ treatment). In contrast, six genes (*PeGATA1, 17, 18, 24, 25* and *26*) showed lower expression in GA_3_-treated seedlings than control seedlings (Fig. 7b). *PeGATA26* was the most down-regulated gene with a 56% expression level reduction, and followed by *PeGATA18* with a 54% decline. It is worth noting that the other 19 *PeGATA* genes did not show significant expression change under GA treatment. These results indicate that the gene expression of *PeGATAs* is at least partially regulated by GA. To test the relationship between *PeGATA* gene expression and auxin, we also analyzed the gene expression pattern of *PeGATA* genes under NAA treatment (5 μΜ NAA) in bamboo seedlings (Fig. 7c, Additional file 6: Table S6). Interestingly, unlike the results of GA treatment, only *PeGATA8* and *PeGATA9* showed significant gene expression change under auxin treatment. Although the expression levels of some other genes including *PeGATA1, 2, 10, 16* and *22* were slightly changed, the gene expression of most *PeGATA* genes did not change under auxin treatment (Fig. 7c), suggesting that PeGATAs may not be affected by auxin. Overall, these results suggest that *PeGATAs* are partially regulated by GA, but are not affected by auxin.

### Genes expression pattern of *PeGATAs* in the rapid-growth of bamboo shoots

As the rapid-growth of bamboo shoots is largely determined by phytohormone and nutrients [2, 27], and we have demonstrated that *PeGATAs* are differentially expressed in rhizome tissues and under GA treatment (Fig. 7a, b), we hypothesized that *PeGATAs* may also be involved in fast-growing bamboo shoots. To validate our hypothesis, the expression profiles of the *PeGATAs* in the fast-growing bamboo shoots (0.15 m, 0.5 m, 1.6 m, 4.2 m and 9 m) were examined by qRT-PCR (Fig. 8). The results showed that almost all GATA genes changed their gene expression in at least one of fast-growing stages. Among them, seven *PeGATA* genes (*PeGATA1, 3, 9, 12, 14, 16, 26* and *31*) continued to decrease their gene expression with the increase of shoot height (Fig. 8). The best example is *PeGATA9*, which showed over 30-fold expression reduction in 9-meter shoots compared to 0.15-meter shoots. The results indicate that these *PeGATAs* genes may be negatively correlated with shoot height. Another groups of *PeGATA* genes (*PeGATA5, 6, 7, 8, 11, 15, 16, 20, 21, 22, 23, 24, 27, 28, 29* and *30*) showed minimal gene expression at the middle growth stages. The results indicate that these *PeGATAs* play an important role in the negative regulation of shoot growth at the middle shoot development stages. Another sets of *PeGATA* genes (*PeGATA2, 4, 10* and *19*) increased their expression at early shoot developmental stages, and then reduce their expression along with the increase of shoot heights. Finally, three *PeGATA* genes (*PeGATA17, 18* and *27*) were increased their expression during early shoot developmental stages, then reduced their expression during middle developmental stages, and then increased their expression again during late shoot developmental stages. Interestingly, we did not find that any *PeGATA* genes continued to increase its expression along with bamboo shoot development. Overall, our results indicate that *PeGATAs* genes may be negatively correlated with rapid-growth of bamboo shoots.

**Fig. 8.**
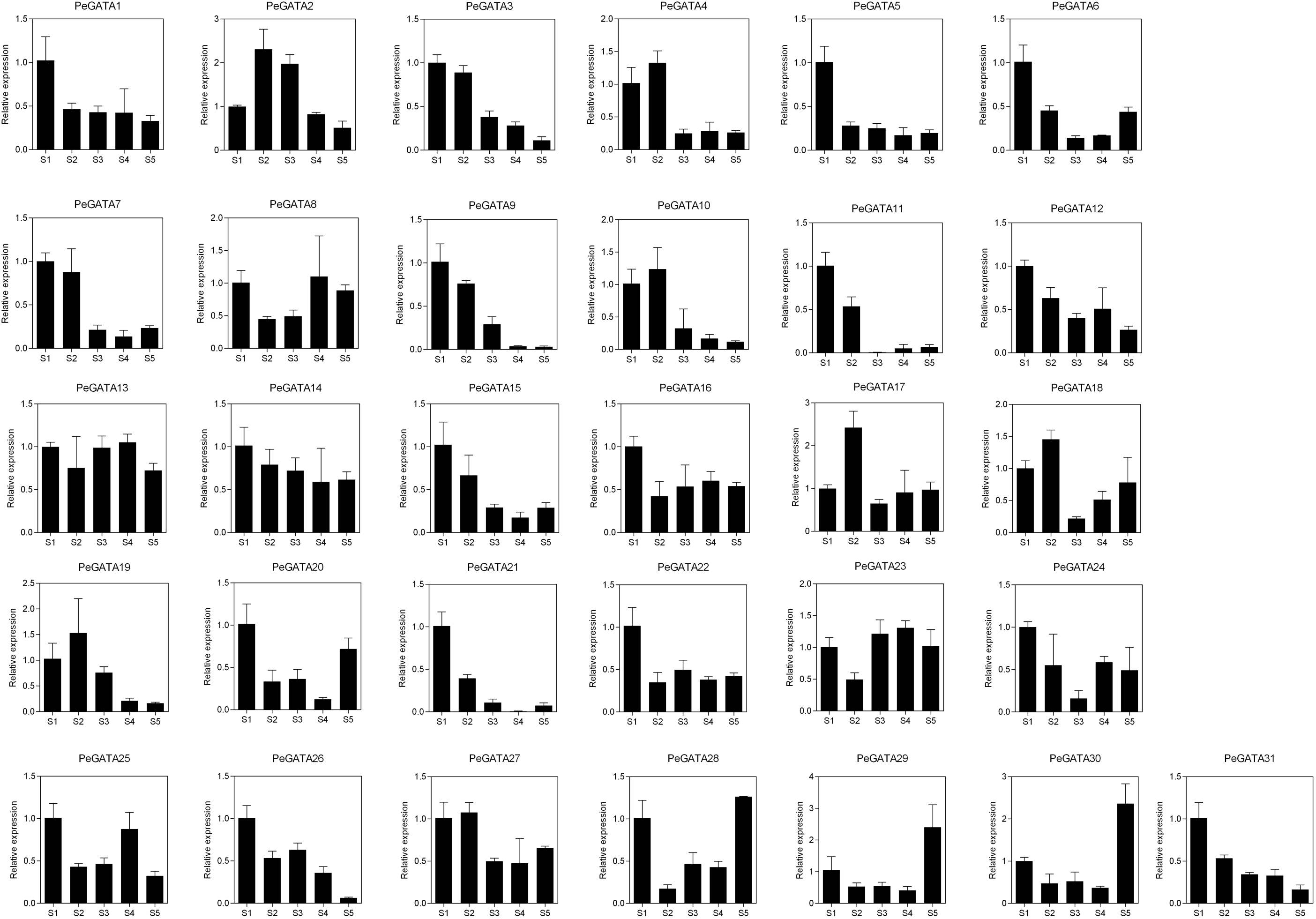
The expression level of *GATA* genes in bamboo shoots. The gene expression values were detected by qRT-PCR. The Y-axis and X-axis indicate relative expression level at different height of shoots. S1: 0.15 m shoots; S2: 0.5 m shoots; S3: 1.6 m shoots; S4: 4.2 m shoots; S5: 9 m shoots.

### Over-expression of *PeGATA26* negatively regulates plant height in Arabidopsis

To understand the function of PeGATA factors, we chose PeGATA26 as an example to verify the role of PeGATA factors in plant growth. *PeGATA26* showed higher gene expression in the growth-inactive lateral buds than the growth-active rhizome tips and new shoot tips (Fig. 7a). Moreover, *PeGATA26* have lower gene expression under GA treatment (Fig. 7b), and its expression decreased along with rapid-growth of bamboo shoots (Fig. 8). These results suggest that *PeGATA26* act as a negative regulator of plant growth and height in moso bamboo. Therefore, we hypothesized that *PeGATA26* plays a crucial role in regulating plant growth. In a previous study, we successfully characterized one of the fast growth-suppressing genes *PeGSK1* by over-expressing it into Arabidopsis [1]. Therefore, we used a similar strategy to verify the function of *PeGATA26* in regulating plant growth by over-expressing it into Arabidopsis.

The homozygous T3 transgenic lines were used to analyze phenotype, and over-expression of *PeGATA26* resulted in a significant growth retardation phenotype (Fig. 9a). Gene expression of *PeGATA26* was successfully detected by qRT-PCR (Fig. 9b). The phenotypes between the two over-expressing lines were similar and the intensity of phenotype correlated with the expression of each transgenic lines (Fig. 9a). Therefore, *PeGATA26* over-expressing line 1 (PeGATA26-ox1) with a stronger phenotype was used for further detailed phenotypic analysis. Interestingly, PeGATA26-ox1 showed a significant dwarf phenotype with a dramatic shorter inflorescence compared to the control (Fig. 9c), indicating that *PeGATA26* inhibits growth of plant height. Moreover, *PeGATA26* also repressed primary root growth (Fig. 9c). However, *PeGATA26* promoted Arabidopsis hypocotyl length (Fig. 9c). These results indicate that *PeGATA26* regulates plant growth in a tissue-specific manner: repressing cell growth in roots and inflorescences, while promoting cell growth in hypocotyls.

**Fig. 9.**
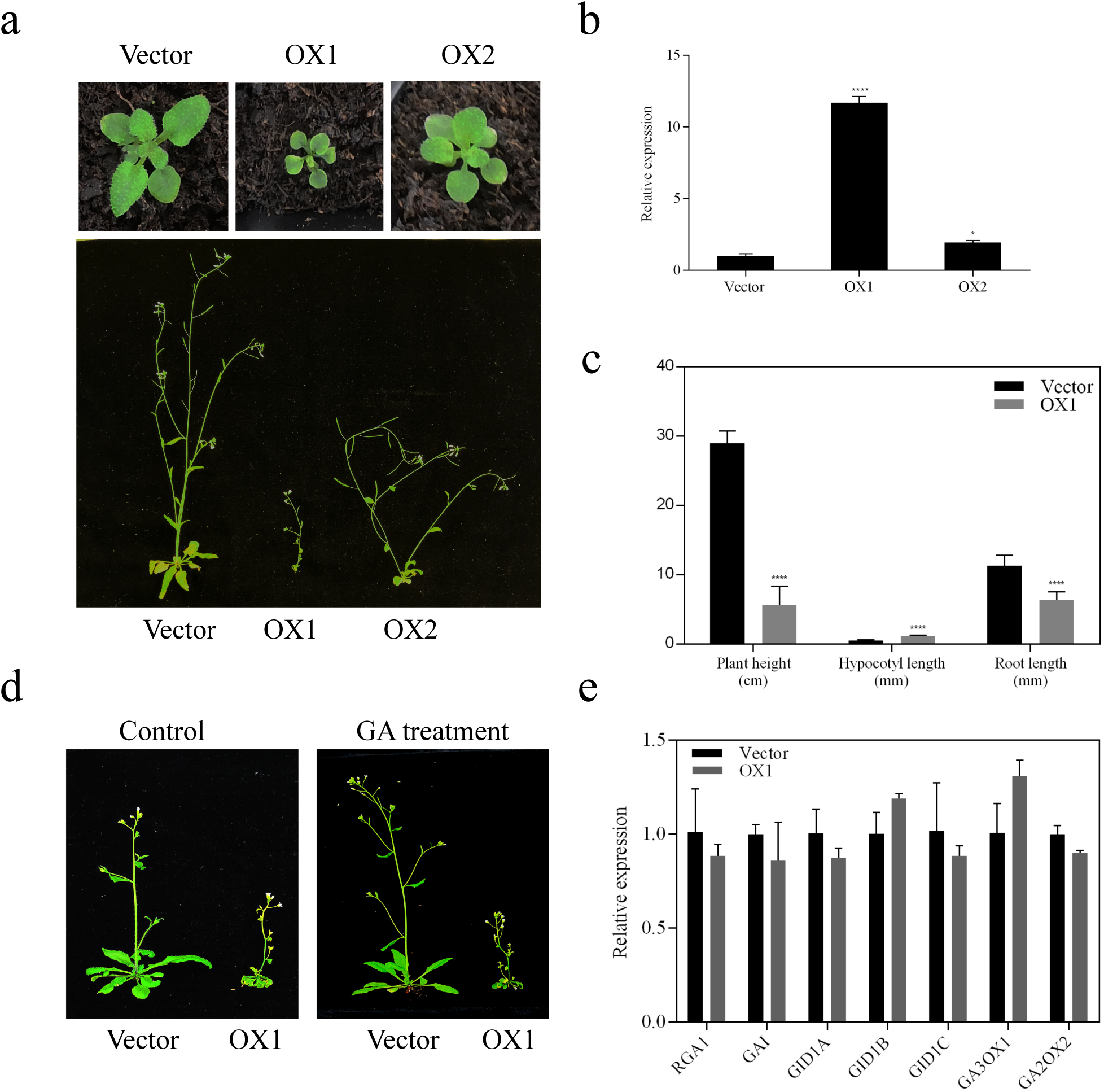
Ectopic expression of *PeGATA26* inhibits the plant height of Arabidopsis. (a): Overexpression of *PeGATA26* resulted in a dwarf phenotype in Arabidopsis. (b): The expression level of *PeGATA26* were detected in both transgenic with the *ACTIN2* gene as an internal control. (c): Phenotypic analysis of plant height, hypocotyl length and primary root length in PeGATA26 overexpressing line 1 compared to control. They all have significant differences compared to the control by the t-test: P ≤ 0.0001, which was represented by four stars in the figure. (d): Overexpression of *PeGATA26* repressed plant height of Arabidopsis and PeGATA26-ox1 was resistant to exogenous GA treatment. (e): The expression of GA signaling genes in the PeGATA26-ox1 was similar to that of GA biosynthesis mutants-*ga1*. The expression of GA signaling genes was detected by qRT-PCR and the *ACTIN2* was used as the internal control.

As *PeGATA26* was down-regulated under the GA treatment in bamboo seedlings (Fig. 7b), we subsequently analyzed whether PeGATA26 was also regulated by GA in Arabidopsis. Interestingly, exogenous GA treatment did not recovery the dwarf phenotype of PeGATA26-ox1 (Fig. 9d), indicating that PeGATA26 is negatively correlated with GA to regulate plant height in Arabidopsis. Similar to the results of its Arabidopsis orthologous gene [21], the gene expression pattern of the GA signaling pathway genes in PeGATA26-ox1 were also similar to those observed in the GA biosynthesis mutants-*ga1* (Fig. 9e). The results support that PeGATA26 also repressed GA signaling downstream of the DELLA protein. These results suggest that PeGATA26 inhibits plant root and stem growth in Arabidopsis, take together with its gene expression negatively correlated with the growth of rhizome tissues and shoot in moso bamboo, we concluded that PeGATA26 is a negative growth regulator for plant height control from Arabidopsis to moso bamboo.

## Discussion

Moso bamboo is one of important non-timber forestry species with great value in providing food and building materials [33]. Moreover, bamboo is known for its fast-growing shoots and widespread rhizomes [2]. It has been reported that several gene families are involved in flower development and abiotic stress [30–32, 41], the rapid-growth associated transcription factors remain elusive. The genome sequences of moso bamboo[33] and transcriptome studies [2, 27, 28, 40] provide important platforms for the identification of rapid-growth shoot and rhizome development associated gene families. The rapid-growth related genes could provide useful information for genetic manipulation of plant height in future.

GATA factors have key functions in developmental control and response to the environmental stresses [16, 17, 19]. In this study, we characterized 31 GATA factors from the moso bamboo (Table 1), and PeGATA factors have highly conserved zinc finger protein domains compared to Arabidopsis and rice GATA factors (Figs. 1-4). More importantly, gene expression analysis of *PeGATAs* in different rhizome tissues and fast-growing bamboo shoots showed that several *PeGATAs* had negative expression patterns with bamboo shoots and rhizome growth (Figs. 7a, 8). Furthermore, the gene expression of *PeGATAs* was partially dependent on phytohormone-GA in bamboo (Figs. 7b). Moreover, overexpression of *PeGATA26* in Arabidopsis repressed the growth of root and plant height in a GA dependent manner (Fig. 9). Overall, our results indicate that *PeGATAs* are involved in regulating the growth of plant height from Arabidopsis to moso bamoo probably through GA signaling pathway.

Bioinformatics analysis showed that there were 31 PeGATA factors in moso bamboo (Fig. 1). The number of bamboo GATA factors was closer to other species, including Arabidopsis (29), rice (28) and apple (35) [9, 10, 15]. Furthermore, most of PeGATA factors have a conserved single CX_2_CX_18-20_ CX_2_C zinc finger domain that is highly similar to that from Arabidopsis and rice (Fig. 1). In addition, the subfamily of I to III from moso bamboo showed a highly evolutionary conservation compared to Arabidopsis and rice (Fig. 3). These results indicate that most of the GATA factors from moso bamboo are conserved compared to other species. However, unlike containing only one zinc-finger domain in subfamily I GATA factors from Arabidopsis and rice [9], PeGATA6 and PeGATA11 from the bamboo GATA subfamily I have two GATA-type zinc finger domains (Fig. 1). Moreover, more protein domains from the bamboo GATA subfamily III were identified compared to that from Arabidopsis and rice (Fig. 2). Interestingly, a unique feature of the PeGATA factors is that they only have three subfamilies compared to the four subfamilies of Arabidopsis and rice (Fig. 3). These differences suggest that PeGATAs do have certain specificity compared to that from Arabidopsis and rice. Future analysis the functions of GATA factors, including AtGATA26, AtGATA27 and OsGATA30 from subfamily IV (Fig. 3), can help us reveal why bamboo lacks these GATA factors.

The first GATA factor is identified according to the light and circadian clock related *cis*-elements in its promoters [13]. Thus, the function of the GATA factors can be predicted based on the identification of *cis*-elements from their promoter. In this study, we found that the promoter of *PeGATAs* has many important *cis*-elements, including light responsive element, cell cycle regulation and phytohormone responsive elements (Fig. 5), which are closely related to the regulation of plant growth. Thus, *PeGATAs* may be involved in regulating plant growth through these *cis*-elements to affect their gene expression and further downstream genes.

The bamboo has a well-established rhizome system to develop new shoot tips and widespread rhizome tips [2, 39]. However, the lateral buds of bamboo rhizomes are not active and dominant in growth [2]. Therefore, identification of GATAs with different expression patterns in these tissues will help us clarify the role of GATA factors in rhizome development, which remains unclear. In this study, we found that 15 *PeGATA* genes are highly expressed in lateral buds (Fig. 7a). Among them, *AtGATA2* (orthologous gene of *PeGATA9)* has been reported to have a function to restrict cell division in the proliferation domain of Arabidopsis root meristem [42], and high expression of *PeGATA9* in lateral buds indicates that *PeGATAs* may also be involved in inhibiting the cell division in bamboo lateral buds (Fig. 7A). In contrast, the lower expression of *PeGATA9* in the actively growing new shoot tips and rhizome tips (Fig. 7A), suggesting a negative correlation between *PeGATA9* and cell growth in bamboo rhizome. It has been reported that *AtGATA22,* a orthologous gene of *PeGATA18,* is involved in response to cytokinin and negatively regulates root growth in Arabidopsis [43], we found that *PeGATA18* has higher gene expression in lateral buds (Fig. 7a), suggesting that *PeGATA18* may play a role in negative regulation of lateral buds cell growth. Similarly, the orthologous gene of *PeGATA26* also plays a negative role in elongation growth [21]. Overall, these results suggest that these 15 *PeGATAs* may contribute to negatively regulating the growth of lateral buds. Next, five *PeGATAs* were highly expressed in the new shoot tips than the other two tissues (Fig. 7a). Among them, the mutation of *GATA19* (orthologous gene of *PeGATA8)* in Arabidopsis causes meristem defects [44]. Here, the high expression of *PeGATA8* in new shoot tips (Fig. 7a), suggests that *PeGATA8* may also be involved in the regulation of shoot meristem development in bamboo. Furthermore, seven *PeGATAs* were highly expressed in the rhizome tips, indicating that they are involved in the growth of the rhizome tips (Fig. 7a). Overall, more than 87% of *PeGATAs* (27/31) was highly expressed in one of rhizome tissues (Fig. 7a). The results indicate that *PeGATAs* strongly participate in the regulation of rhizome growth. Once the transformation system is ready in future, functional characterization of these *PeGATAs* in moso bamboo will help us elucidate the exact role of PeGATAs in rhizome growth control.

The correlation between GATA factors and GA or auxin has been extensively studied in Arabidopsis [21, 23, 42, 45]. In this study, we found that gene expression of 12 *PeGATAs* changed under GA treatment, while only two *PeGATAs* responded to auxin treatment (Fig. 7b, c). In addition, motif analysis indicated that the promoter of *PeGATAs* has more GA-related *cis*-elements than auxin (Fig. 5b). Our results indicate that GA rather than auxin frequently regulates the expression of *PeGATAs* in moso bamboo.

Gene expression analysis showed that almost all PeGATAs have changed their expression during the rapid-growth of bamboo shoots (Fig. 8). For example, *GATA2* (orthologous gene of *PeGATA9*) limits cell division in root meristem of Arabidopsis [42], and the expression of *PeGATA9* was down-regulated over 30 times in late rapid-growth stage (9 m) than the early stage (0.15 m) (Fig. 8). The results indicate that *PeGATA9* may also negatively regulate the rapid-growth of bamboo shoot. Identification of many rapid-growth related *PeGATAs* indicates that *PeGATAs* are involved in regulating the bamboo shoot. The rapid-growth of bamboo shoot is tightly controlled by phytohormone [27]. Current studies reveals that ABA is the only negative regulator of fast-growing shoots, while BR, auxin, GA and cytokinin antagonize with ABA to promote rapid-growth of bamboo shoots [27]. Interestingly, all of the 12 GA-related *PeGATAs* showed differential expression in at least one of rapid-growth stages (Figs. 7b, 8), suggesting that GA may regulate rapid-growth of bamboo shoots via modulating gene expression of *PeGATAs*. To understand the function of GA-related *PeGATAs* in plant height control, *PeGATA26* was selected to validate its role in Arabidopsis growth (Fig. 8). Overexpression of *PeGATA26* in Arabidopsis resulted in growth retardation phenotypes such as dwarfism and shorter primary root length, and the *PeGATAs* over-expressed lines was resistant to GA treatment (Fig. 8). Overall, these results further support that *PeGATAs* could regulate plant heights from Arabidopsis to moso bamboo via GA signaling pathway.

### Conclusions

With the explosive growth rates of bamboo shoots and widespread rhizomes, the identification of key regulatory genes in the bamboo shoot and rhizome growth control will provide important genetic resources for the genetic manipulation of plant height. In this study, we characterized 31 GATA factors from moso bamboo. More importantly, the gene expression of *PeGATAs* is closely related to the development of rhizome tissues and rapid-growth of bamboo shoots. Moreover, the gene expression of *PeGATAs* was partially regulated by the phytohormone-GA in bamboo. In addition, functional characterization of *PeGATA26* in Arabidopsis provides insight into how *PeGATAs* regulate plant height from Arabidopsis to bamboo via the GA signaling pathway. However, we also noticed that GA regulates expression of only part of PeGATAs. As ABA-related *cis*-elements are more widespread than GA, and ABA is the only known negative regulatory hormones in the rapid-growth control of bamboo shoots, we cannot rule out that *PeGATAs* may also regulate plant height through ABA signaling pathway. In summary, our results provide certain evidence that GATA transcription factor regulate the development of rhizome tissues and the rapid-growth of bamboo shoots.

## Methods

### Identification of GATA factors in moso bamboo

To identify the GATA factors, the genome and protein sequences of moso bamboo were downloaded from BambooGDB database (http://forestry.fafu.edu.cn/db/PhePacBio/download.php) [33]. GATA protein sequences from Arabidopsis and rice were obtained from previous published data [9]. We performed multiple sequence blast and alignment with an expected value of 10. The HMMER profile of the GATA domain (PF00320) from Pfam (http://pfam.xfam.org/) was used to search the bamboo protein database with a threshold: e-values < 10^-5^ [46]. The bamboo GATA factors were determined with the criteria: it is present in both blast and HMMER motif analysis lists. The number of amino acid, molecular weights (MWs) and isoelectric points (PI) of bamboo GATA factors were predicted by ProtParam (https://web.expasy.org/protparam/).

### Phylogenetic tree, conserved domain, motif recognition and *cis*-elements analysis

Multi-sequence alignment of the GATA protein sequences was carried out by ClustalX [47], and phylogenetic tree was constructed using MEGA7 by the Neighbour-Joining method (bootstrap analysis for 1000 replicates) [48]. Conserved domains were obtained from NCBI (https://www.ncbi.nlm.nih.gov/cdd) [49] and motifs were analyzed using MEME with default parameters (version 5.0.5, http://meme-suite.org/tools/meme) [50]. For *cis*-elements analysis, DNA sequences from 1.5-kb upstream region of each *PeGATA* gene were used to scan any potential *cis*-element using the PlantCARE database (http://bioinformatics.psb.ugent.be/webtools/plantcare/html/) [51].

### Subcellular localization analysis

To verify the location of PeGATAs, the full-length CDSs without stop codon from four *PeGATA* genes were cloned into a modified pCambia3301 vector with C-terminal GFP as described in our previous study [1]. The *ACTIN2::PeGATAs*-GFP and the *ACTIN2*::GFP control constructs were then transiently transformed into tobaccos, and GFP and DAPI fluorescence was observed using a microscope (20x, Zeiss, LSM880).

### Gene expression analysis

To investigate gene expression levels of the PeGATA genes in different tissues or hormone treatments, RNA-seq data was downloaded from Short Read Archive (SRA) database for the lateral buds, rhizome tips and new shoot tips (SRP093919) [2], and bamboo seedlings under GA and auxin treatment (SRP119416 and SRP109631) [28, 40], respectively. The pair-end reads were mapped to the moso bamboo reference genome using tophat2, and differential expressed genes were detected by cufflinks with defaults parameters [52].

### Plant materials and qRT-PCR analysis

The moso bamboo shoots used in this study were collected in JianOu County (E118°28’; N27°00’), Fujian Province, China. The middle internode of different height of bamboo shoots were sampled and stored in liquid nitrogen immediately. qRT-PCR analysis was performed for each member of the GATA family genes during the rapid-growth of bamboo shoots. Total RNA was extracted from the bamboo samples using HiPure Plant RNA Mini Kit (Magen, R4151-02) and 1μg RNA was taken for reverse transcription into cDNA using a commercial Kit (Monad, RN05004M). Primers for qRT-PCR were designed on Primer3 (http://primer3.ut.ee/) using the CDS of each PeGATA gene. qRT-PCR were performed using MonAmp™ ChemoHS qPCR Mix (Monad, RN04002M) in a 20 μl reaction. The following program was used for qRT-PCR: 95 °C for 5 min; 40 cycles of 95 °C for 10 s, 60 °C for 10 s and 72 °C for 30 s.

### Ectopic expression analysis

The *PeGATA26* was cloned and expressed in Arabidopsis exactly following the procedures for the *PeGSK1* in our previous study[1]. The T3 generation seedlings were used for phenotype analysis. The primary root length, plant height and hypocotyl length were measured using ImageJ[1].

The primers used in this study were listed in Additional file 7: Table S7.

## Declarations

### Competing interests

The authors declare that they have no competing interests.

### Funding

This work was supported by the National Natural Science Foundation of China (Nos. 31741025 and 31500258 to L.M), the Outstanding Youth Research Talents Development Program in Fujian Province University to Liuyin Ma, the Outstanding Youth Research Talents Program of Fujian Agriculture and Forestry University (No. KXJQ17011 to L.M.), and the Scientific Research Foundation of the Graduate School of Fujian Agriculture and Forestry University (to T.W.).

### Author contributions

T.W., C.L., and L.M. conceived the ideas. T.W. and W.W. performed the experiments. T.W., Y.Y., S.L. and Z.Z. contributed to data analysis. T.W. and L.M. wrote the manuscript.

## SUPPORTING INFORMATION

### Additional files

**Additional file 1: Table S1.** The coding region sequences of *PeGATA* genes.

**Additional file 2: Table S2.** The amino acid sequences of PeGATA factors.

**Additional file 3: Table S3.** List of protein motifs identified in PeGATA factors.

**Additional file 4: Table S4.** Gene expression of *PeGATA* genes in different rhizome tissues.

**Additional file 5: Table S5.** Gene expression of *PeGATA* genes under GA treatment.

**Additional file 6: Table S6.** Gene expression of *PeGATA* genes under auxin treatment.

**Additional file 7: Table S7.** List of primers used in this study.

